# Functional and transcriptional profiling of non-coding RNAs in yeast reveal context-dependent phenotypes and widespread *in trans* effects on the protein regulatory network

**DOI:** 10.1101/2020.04.07.029611

**Authors:** Laura Natalia Balarezo-Cisneros, Steven Parker, Marcin G Fraczek, Soukaina Timouma, Ping Wang, Raymond T O’Keefe, Catherine B Millar, Daniela Delneri

## Abstract

Non-coding RNAs (ncRNAs), including the more recently identified Stable Unannotated Transcripts (SUTs) and Cryptic Unstable Transcripts (CUTs), are increasingly being shown to play pivotal roles in the transcriptional and post-transcriptional regulation of genes in eukaryotes. Here, we carried out a large-scale screening of ncRNAs in *Saccharomyces cerevisiae*, and provide evidence for SUT and CUT function. Phenotypic data on 372 ncRNA deletion strains in 23 different growth conditions were collected, identifying ncRNAs responsible for significant cellular fitness changes. Transcriptome profiles were assembled for 18 haploid ncRNA deletion mutants and 2 essential ncRNA heterozygous deletants. Guided by the resulting RNA-seq data we analysed the genome-wide dysregulation of protein coding genes and non-coding transcripts. Novel functional ncRNAs, SUT125, SUT126, SUT035 and SUT532 that act *in trans* by modulating transcription factors were identified. Furthermore, we described the impact of SUTs and CUTs in modulating coding gene expression in response of different environmental conditions, regulating important biological process such as respiration (SUT125, SUT126, SUT035, SUT432), steroid biosynthesis (CUT494, SUT530, SUT468) or rRNA processing (SUT075 and snR30). Overall, this data captures and integrates the regulatory and phenotypic network of ncRNAs and protein coding genes, providing genome-wide evidence of the impact of ncRNAs on cellular homeostasis.

**Author Summary:** The yeast genome contains 25% of non-coding RNA molecules (ncRNAs), which do not translate into proteins but are involved in regulation of gene expression. ncRNAs can affect nearby genes by physically interfering with their transcription (*cis* mode of action), or they interact with DNA, proteins or others RNAs to regulate the expression of distant genes (*trans* mode of action). Examples of *cis*-acting ncRNAs have been broadly described, however genome-wide studies to identify functional *trans*-acting ncRNAs involved in global gene regulation are still lacking. Here, we used the ncRNA yeast deletion collection to score their impact on cellular function in different environmental conditions. A group of 20 ncRNAs mutants with broad fitness diversity were selected to investigate their effect on the protein and ncRNA expression network. We showed a high correlation between altered phenotypes and global transcriptional changes, in an environmental dependent manner. We confirmed the widespread *trans* acting expressional regulation of ncRNAs in the genome and their role in affecting transcription factors. These findings support the notion of the involvement on ncRNAs in fine tuning the cellular expression via regulations of TFs, as an advantageous RNA-mediated mechanism that can be fast and cost-effective for the cells.

## Introduction

Gene regulation is a key biological process across all life forms, and multiple gene interactions quickly allow adaptation to different conditions in response to environmental stimuli. This response may induce adaptation to various food sources, trigger alternative metabolic pathways, or overcome stress factors.

Chromatin modifications and DNA methylation are two main mechanisms of regulating gene expression. More recently, RNA transcripts which are not translated into protein, have been described to have a prominent role as epigenetic modifiers [1,2]. There are an increasing number of examples of these non-coding RNA (ncRNA) transcripts regulating gene expression positively and negatively [3-10].

RNA interference (RNAi) was the first understood example of ncRNA involvement in epigenetics [11]. This RNAi mechanism involves ncRNAs binding to target mRNA sequences, inhibiting their translation [12]. *Saccharomyces cerevisiae* (*S. cerevisiae*) lacks RNAi machinery; however, a large number of non-coding transcripts have been identified in this budding yeast using high-throughput and high-resolution technologies. These ncRNA transcripts come from what is known as “pervasive transcription”, a mechanism that generates RNAs distinct from those that encode proteins or those with established functions (*e.g*. snoRNAs, snRNAs, rRNAs) [13]. Among a list of catheterized pervasive transcripts, Stable Unannotated Transcripts (SUTs) and Cryptic Unstable Transcripts (CUTs) show an essential role in gene regulation, influencing histone modifications or regulating transcription of nearby genes [4,5,14–16].

SUTs and CUTs are polyadenylated RNAs transcribed by RNA polymerase II [17] and are distributed across the entire *S. cerevisiae* genome. Classically, SUTs and CUTs arise from nucleosome-depleted regions (NDRs) associated with bidirectional promoters of protein-coding genes [17,18], but differ in their association with the RNA decay machinery. CUTs are capped and degraded rapidly by the nuclear exosome and the TRAMP (Trf4-Air1/Air2-Mtr4) complex [19], whereas SUTs are only partially susceptible to Rrp6p activity [17] and are mainly affected by cytoplasmic RNA decay pathways including the translation-dependent nonsense-mediated decay (NMD) pathway and Xrn1-dependent 5’ to 3’ degradation [20]. As a result, SUTs persist longer than CUTs.

Gene regulation activities have been ascribed to SUTs and CUTs. In many cases, ncRNAs appear to cause transcriptional interference [3,5,15,16,21-23] affecting the expression of neighbouring genes in *cis*. On the other hand, ncRNAs can be functional and regulate in *trans* the expression of genes located both nearby or at distant loci [4,6,10]. Although only a small number of functional ncRNAs have been well characterized to date, they have been shown to control gene expression at the transcriptional level. For instance, SUT075 has recently been reported to regulate the expression of *PRP3* when overexpressed remotely on a plasmid [10]. Another example is SUT457, which is involved in telomere organization. SUT457 regulates the levels of telomeric ssDNA in a Exo1-dependent manner [9]. Interestingly, CUT281, known as *PHO84* ncRNA because it overlaps the protein-coding *PHO84* gene, triggers *PHO84* silencing in a *trans* and *cis* manner using two independent mechanisms. While the *cis-*acting mechanism requires Hda1/2/3 deacetylation machinery, *trans* function is generated by the Set1 histone methyltransferase [5,6].

Emerging evidence has suggested ncRNAs roles in the recruitment of transcription factors (TFs) to their binding sites in fission yeast, mouse and humans [21-26], thus, suggesting a conserved mechanism of gene expression among eukaryotes. On one hand, ncRNA expression around regulatory elements can locally promote TF binding [23-24]. On the other hand, ncRNA can regulate gene expression by acting as binding competitors for DNA-binding proteins (DBPs) [25-26].

Considerable progress has been made over the past decade to elucidate the unique features and molecular mechanisms of ncRNA. However, detailed insights have been limited to single ncRNA genes, usually affecting neighbouring genes. Here, we combine large-scale phenotypic analysis with RNA-seq technology to generate a global view of the transcriptome following ncRNA deletion. Specifically, by analysing the expression network, we show that the global transcriptional effect of deleting four SUTs is indirect and acts via specific TFs whose level of expression is affected by deleting these ncRNAs. This *trans* effect supports and extends previous premises that SUTs or CUTs are a functional part of the genome and can influence the general transcriptional output of a cell independent from where they are located.

## Results and Discussion

### Fitness profiling of haploid ncRNA deletion strains reveals plasticity of phenotype in different environmental conditions

To investigate the plasticity of ncRNA deletion mutations on organism fitness, we acquired phenotypic data for the haploid ncRNA deletion collection generated by Parker *et al*. (2017) in 23 different conditions. The ability of 50 CUT, 93 SUT, 61 snoRNA and 168 tRNA deletion mutant strains to utilize different carbon sources, and to tolerate extreme pH and oxidative stress was scored. The colony size was used as a proxy for fitness and normalized to the wild-type strain according to Tong and Boone [9]. The ncRNA deletion mutants showing similar behaviour across the 23 different conditions were grouped, generating 42 distinct functional clusters (Fig 1). The list of deletion mutant strains in each cluster is reported in the Supplementary Dataset S1.

**Fig 1.**
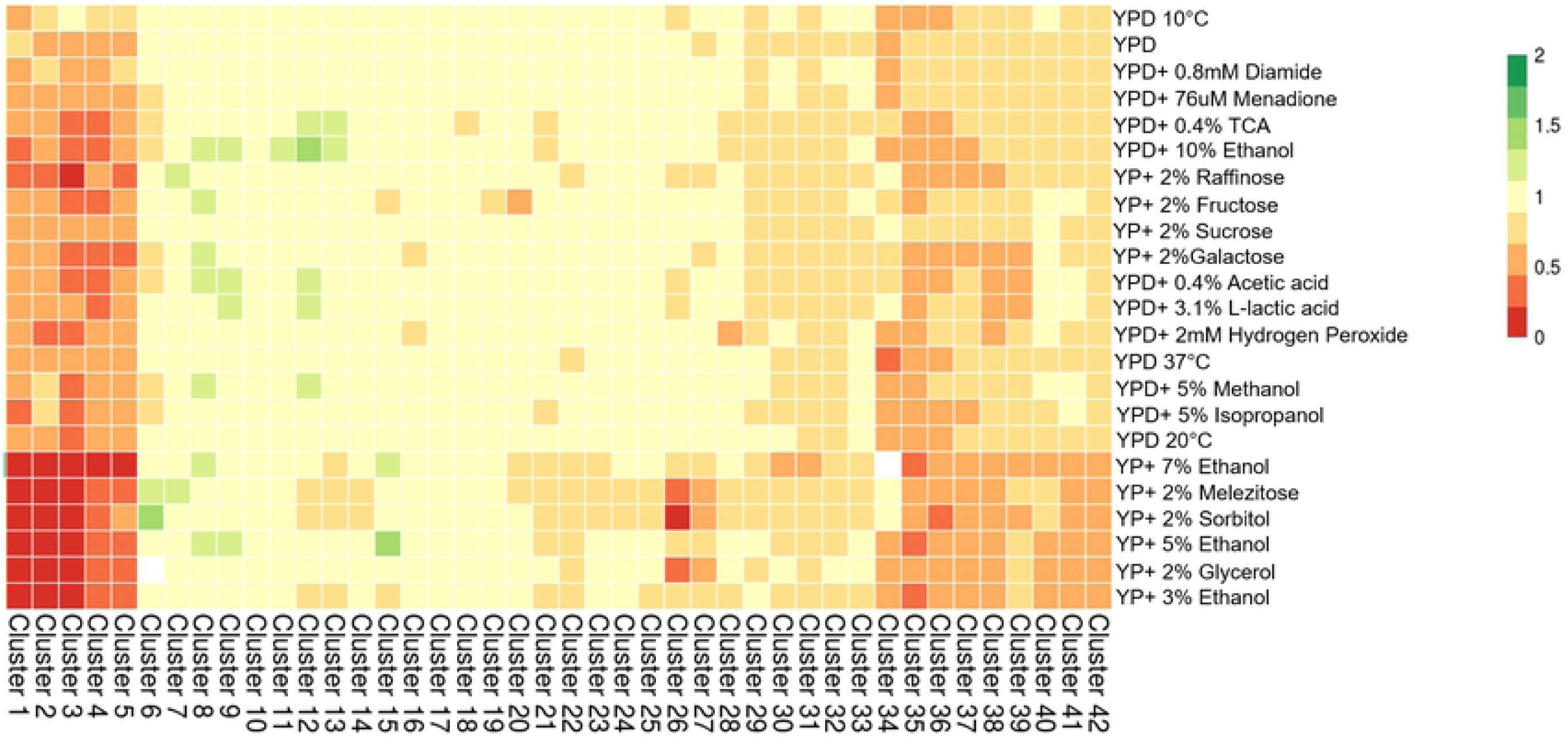
ncRNA deletion strain fitness profiles on solid media in different environments. Heat-map of the 42 clusters containing 372 haploid ncRNA deletion strains. Rows represent the different growth conditions and columns represent the clusters. Colour bars represent the colony size normalized to the wild-type strain which is given the arbitrary growth value of 1. Fitness reduction is represented as shades of red. Fitness increased is represented as shades of green. No fitness change is represented as yellow. Missing data is represented as white. The list of deletion mutant strains in each cluster can be found in Supporting information S1 Dataset.

About 45% of the ncRNA deletion mutants analysed did not show significant phenotypic changes in any condition tested, while about 24% of the ncRNA deletion strains showed significant changes in fitness in at least one condition tested. The remaining ncRNA deletion mutants (*i.e*. clusters 1 to 5) displayed a severe fitness defect in the majority of conditions, in particular when grown in ethanol, glycerol, sorbitol or melezitose as carbon sources. These clusters contained mostly tRNAs and SUTs rather than CUTs and snoRNAs (Supplementary Dataset S1). The conditions that affected the least number of ncRNA deletion mutants were YP+ 2% Fructose, YPD+ 5% Methanol, and YPD+ 5% Isopropanol, which affected 4.3%, 4.5% and 5.1% of ncRNA deletion mutants, respectively. Conditions that induced the broadest fitness changes were YP+ 7% Ethanol and YP+ 2% Glycerol, with 11.8% and 11.5% of ncRNA deletion mutants affected, respectively (Fig 1 and Supplementary Dataset S1).

Liquid growth assays were also set up for SUT and CUT deletion mutants that displayed either severe fitness defects (clusters 1, 2 and 5), fitness gain (clusters 7 to 9), or no phenotype (cluster 10). The overall liquid growth phenotype is reported as relative area under the growth curve (Fig 2), and the breakdown for the different growth phases is available in Table S1. Overall, in rich media, the majority of deletion mutant strains showed no growth difference, with the exception of the reduced fitness of *SUT125*Δ, *SUT126*Δ and CUT494/SUT053/SUT468*Δ* and the improved fitness of *CUT248Δ* (S1 Table).

**Fig 2.**
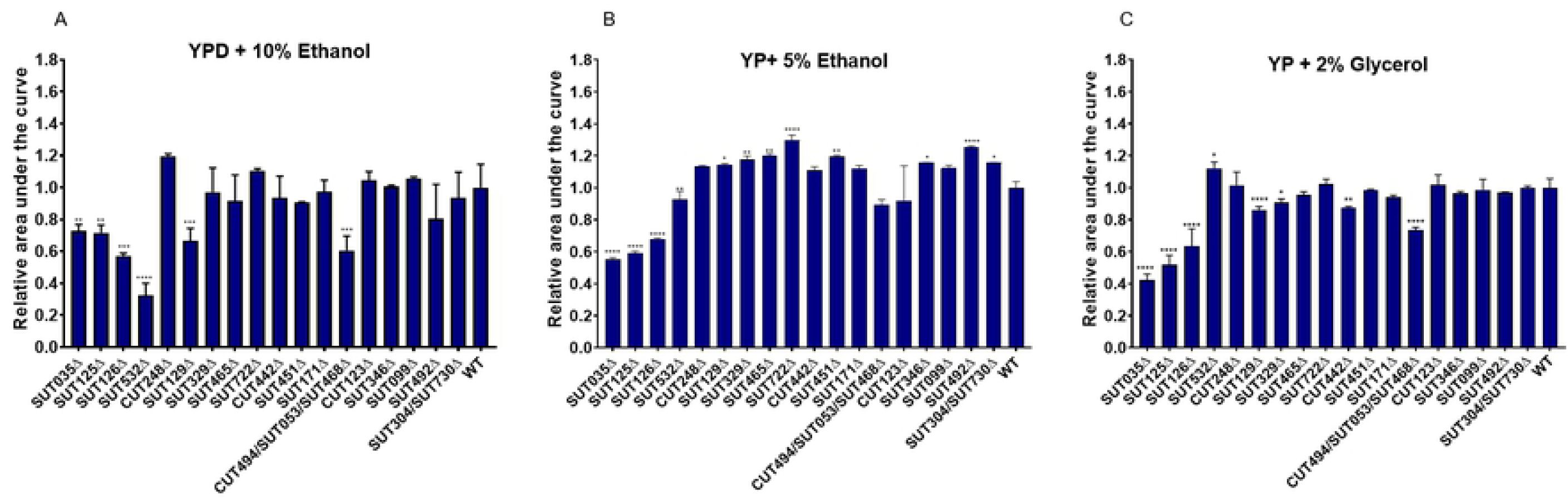
Liquid growth assays for SUT and CUT deletants with pronounced growth differences on solid media. Bar charts show the relative area under the curve for haploid SUT and CUT deletion strains grown in (A) YPD+10% Ethanol (B) YP+ 5% Ethanol and (C) YP+2% Glycerol. The data are presented as means calculated from three biological replicates normalized to WT. Comparisons between wild-type and mutants were analysed using ANOVA followed by Dunnett’s test.

When 10% ethanol was added to the media, *SUT125Δ, SUT126Δ, SUT035Δ, SUT532Δ, SUT129Δ* and *CUT494/SUT053/SUT468Δ* displayed severe fitness defects affecting the majority of the growth phases (Fig 2A and S1 Table); a similar profile for fitness impairment for SUT125, SUT126 and SUT035 was observed in media containing either 5% ethanol (Fig 2B) or 2% glycerol (Fig 2C). However, *SUT532Δ* and *SUT129Δ* had a divergent fitness profile in 5% ethanol and 2% glycerol. *SUT532Δ* presented a significant fitness defect in YP+ 5% Ethanol and a growth improvement in YP+ 2% Glycerol (Fig 2B and C), whereas *SUT129Δ* showed an improvement in YP+5% Ethanol and defect in 2% glycerol.

Several SUTs and CUTs displayed improved fitness in the YP+ 5% Ethanol liquid media (Fig 2B) revealing a similar phenotypic change in both solid and in liquid media. About 56% of the strains grown in YPD+10% Ethanol and 27.7% of the strains grown in YP+ 2% Glycerol displayed some differences in fitness profiles between solid and liquid media. For example, *CUT494/SUT053/SUT468Δ, SUT129Δ, SUT329Δ* and *CUT442Δ* exhibited fitness impairment in YP+ 2% Glycerol which was not previously detected in the solid fitness assay. Discrepancies between solid and liquid fitness are likely due to the differing oxygen availability and diffusion rates of one or more nutrients on solid media [27-31]. Indeed, when growing on solid surfaces, colony morphology differs between yeast growth phases and time [32–34]. Therefore, these results re-iterate the importance of acquiring data from both solid and liquid growth assays for an accurate picture of cellular fitness.

### ncRNA deletions drive global transcriptional changes that correlate with phenotypic profiles

The main function previously ascribed to ncRNAs in budding yeast is transcriptional regulation, usually of neighbouring or overlapping single genes [3, 4, 7, 16, 35, 36] We therefore investigated by RNA-seq whether selected ncRNA deletion mutants with altered phenotypes also have dysregulated transcriptomes. We selected 18 haploid ncRNA deletion mutants from clusters 1, 2, 5 and 7-10 with different types of phenotypic changes (*i.e*. growth defects, improvements and no changes) to study by RNA-seq, together with heterozygous deletions of 2 essential ncRNAs, namely SUT075 and snR30 (previously described in Parker et al [10]) (Table 1). As expected, we detected changes in the levels of at least one neighbouring transcript in 8 of the ncRNA deletion mutant strains analysed by RNA-seq. Three of these deletion mutants (*SUT099Δ, SUT722Δ, SUT171Δ*) up-regulate only their neighbouring genes, while the remainder (*CUT494/SUT053/SUT468Δ, SUT532Δ, SUT035Δ*, and *SUT125Δ*) also revealed altered levels of distantly located transcripts. Strikingly, over one-third of the deletion mutants studied by RNA-seq had large numbers (>100) of differentially expressed (DE) coding and non-coding transcripts (Table 1).

**Table 1.** Numbers of protein-coding genes and non-coding transcripts that are differentially expressed in 18 SUT and CUT deletants. A ‘neighbour’ gene or transcript is defined as an adjacent genomic feature.

| Protein- coding genes |  |  |  |  |  | Non-coding DNA |  |  |  |
| --- | --- | --- | --- | --- | --- | --- | --- | --- | --- |
|  | SUT/CUT | Number of DE genes | Up - Regulated | Down- Regulated | Neighbour DE gene | Number of DE transcripts | Up - Regulated | Down- Regulated | Neighbour DE transcript |
| **Cluster 1** | SUT035 | 701 | 256 | 445 | 1 | 232 | 138 | 94 | 0 |
|  | SUT125 | 721 | 310 | 411 | 2 | 196 | 106 | 90 | 0 |
| **Cluster 2** | SUT126 | 787 | 335 | 452 | 0 | 223 | 141 | 82 | 0 |
| **Cluster 5** | SUT532 | 408 | 236 | 172 | 0 | 92 | 55 | 37 | 1 |
| **Cluster 7** | CUT494/<br>SUT530/<br>SUT468 | 137 | 102 | 35 | 1 | 32 | 6 | 26 | 0 |
|  | CUT123 | 0 | 0 | 0 | 0 | 0 | 0 | 0 | 0 |
|  | CUT248 | 2 | 2 | 0 | 0 | 0 | 0 | 0 | 0 |
| **Cluster 8** | SUT129 | 16 | 8 | 8 | 0 | 2 | 1 | 1 | 0 |
|  | SUT304/<br>SUT730 | 1 | 0 | 1 | 0 | 0 | 0 | 0 | 0 |
|  | SUT329 | 1 | 0 | 1 | 0 | 0 | 0 | 0 | 0 |
|  | SUT722 | 1 | 1 | 0 | 1 | 0 | 0 | 0 | 0 |
| **Cluster 9** | SUT171 | 2 | 1 | 1 | 1 | 0 | 0 | 0 | 0 |
|  | SUT346 | 1 | 0 | 1 | 0 | 0 | 0 | 0 | 0 |
|  | SUT465 | 0 | 0 | 0 | 0 | 0 | 0 | 0 | 0 |
| **Cluster 10** | CUT442 | 2 | 0 | 2 | 0 | 0 | 0 | 0 | 0 |
|  | SUT099 | 2 | 1 | 1 | 1 | 0 | 0 | 0 | 0 |
|  | SUT451 | 0 | 0 | 0 | 0 | 0 | 0 | 0 | 0 |
|  | SUT492 | 1 | 0 | 1 | 0 | 0 | 0 | 0 | 0 |

Half of the deletion mutant strains had smaller numbers of differentially expressed transcripts, while only three deletion mutants did not lead to transcriptional changes in rich medium. Overall, transcription profiles of the ncRNA deletion mutants correlated well with their fitness changes. For instance, heterozygous deletions of the two essential ncRNAs SUT075 and snR30 have overall a stronger negative effect on strain fitness in all the conditions tested (S1 Fig). As expected, these deletions affected the largest number of transcripts (Table 2).

**Table 2.** Differentially expressed protein-coding genes and non-coding transcripts in heterozygous deletions of the essential ncRNA genes *snR30* and *SUT075*.

|  | Protein-coding genes |  |  |  | Non-coding DNA |  |  |  |
| --- | --- | --- | --- | --- | --- | --- | --- | --- |
|  | Number of DE genes | Up - Regulated | Down-Regulated | Neighbour | Number of DE transcripts | Up - Regulated | Down-Regulated | Neighbour |
| snR30 | 2276 | 1063 | 1213 | 0 | 408 | 335 | 73 | 2 |
| SUT075 | 2284 | 1057 | 1227 | 1 | 292 | 238 | 54 | 0 |

The two apparently unrelated essential ncRNAs, SUT075 and snR30, have a surprisingly large number of DE transcripts in common (about 80%; 864 up-regulated and 972 down-regulated). Gene Ontology (GO) analysis of the shared DE protein-coding genes revealed enrichment for ribosome biogenesis, ribosomal RNA processing, DNA replication and the cell cycle (S2 Fig). This GO enrichment is consistent with the known role of snR30 in ribosomal RNA processing [37]. SUT075 is required for normal transcript levels of its neighbouring essential gene *PRP3* and can act in *trans* [10]. We note however, in our RNA-seq data, that the down-regulation of *PRP3* was not significantly strong (fold change, FC, 0.7) in *SUT075*Δ and instead a large global effect on the transcriptome was detected, including targets in common with the snR30 mutant. A further 481 essential genes are affected in addition to *PRP3* when SUT075 is deleted (82 up-regulated and 399 down-regulated) representing 43% of the *S. cerevisiae* essential genes. As a comparison, *snR30*Δ dysregulates 450 essential genes (ca. 40%), up-regulating and down-regulating 82 and 368, respectively. Nineteen small RNAs are dysregulated in *snR30*Δ, including the essential snR19 and LSR1 (U1 and U2 snRNAs) that are part of the major spliceosome in yeast, and the RNA component of nuclear RNase P (*RPR1*) and RNase MRP (*NME1*). Interestingly, 17 snoRNAs, of which 11 are in common with *snR30*Δ, are also differentially expressed in *SUT075*Δ. Our data suggest that several factors, including an effect on the neighbouring gene *PRP3* and a potential role in rRNA processing, may cause the essentiality of SUT075.

### Discordant changes between transcriptome and fitness as a tool to reveal additional context-dependent phenotypes

Generally, the clusters containing ncRNA deletion mutants that do not impair fitness also do not produce significant global transcriptional changes and vice-versa. However, there was one exception to this pattern in cluster 7 that contained ncRNA deletion mutants with largely unaffected fitness except for *CUT494/SUT053/SUT468Δ*. The deletion of CUT494/SUT053/SUT468 caused the dysregulation of over 150 transcripts, in contrast to the deletion of CUT123 or CUT248, which affected 0 and 2 transcripts respectively. To investigate this apparent discrepancy, we used GO analysis to identify enriched functional categories across the 137 DE genes in *CUT494/SUT053/SUT468Δ*. The majority of the over-represented GO terms were related to the synthesis of crucial membrane components and membrane fluidity pathways. Specifically, GO biological process categories enriched among down-regulated genes included sterol, steroid, ergosterol and lipid biosynthesis, while up-regulated genes were clustered in pathways for propionate metabolism, drug response and molecular transport (S3 Fig). Ergosterol (ERG) is an essential membrane component that regulates membrane fluidity, permeability, membrane-bound enzyme activity and substance transportation [38]. Overexpression or deletion of ERG biosynthesis genes results in the accumulation of toxic intermediates, alteration of drug sensitivity and slow growth in different media, including non-fermentable carbon sources [39]. Interestingly, our fitness data revealed a growth defect of the *CUT494/SUT053/SUT468*Δ strain in YP+2% Glycerol (Fig 2C). To test the hypothesis that *CUT494/SUT053/SUT468*Δ has a role in membrane stability by targeting synthesis of ERG, we used azole antifungal agents that inhibit various steps in the ERG biosynthesis pathway [40]. When the fitness of *CUT494/SUT053/SUT468Δ* was tested in medium supplemented with either fluconazole (Fig 3A and C) or miconazole (Fig 3B and D), a slow growth phenotype was identified compared to the WT and the other deletion mutants in the same cluster (Fig 3). This result suggests that transcriptome data can be used to identify environmental conditions that are likely to reveal fitness defects.

**Fig 3.**
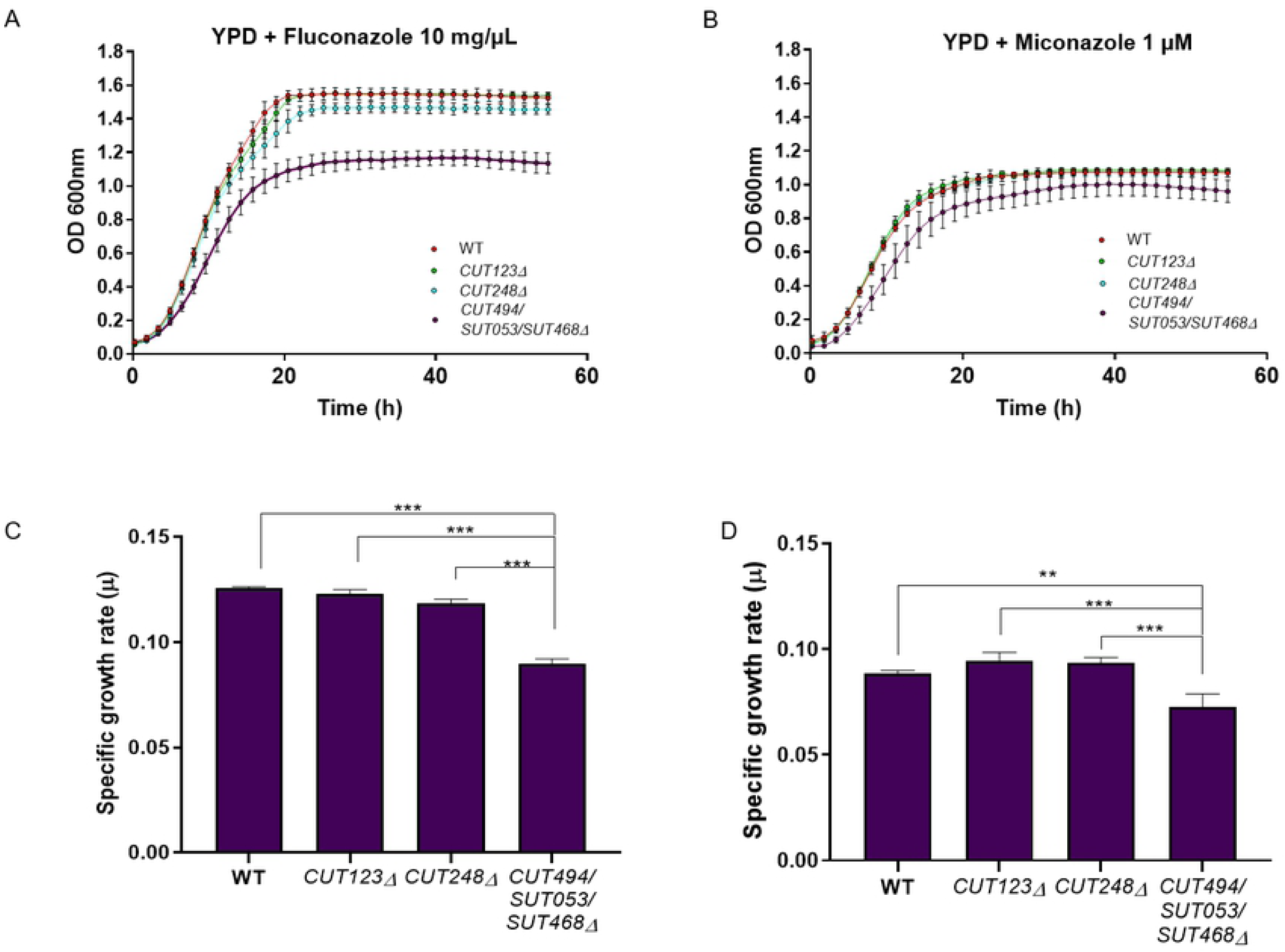
Liquid growth assays of ncRNA deletion mutants in the presence of azoles. Growth curves of *CUT123Δ, CUT248Δ, CUT494/SUT530/SUT468Δ* and WT strains in YPD media supplemented with (A) Fluconazole (10 mg/μl) and (B) Miconazole (1 µM). Bar charts show the mean specific growth rate (μ) of WT and ncRNA deletion strains in the presence of (C) fluconazole (10 mg/μl) and D) miconazole (1 μM). Significance of differences was assessed by *t*-tests.

### ncRNAs with related phenotypes regulate common genes involved in mitochondrial functions

SUTs/CUTs clustered together by their fitness profile are expected to engage similar biological and molecular functions. To test this premise, we identified the set of common DE genes across deletion mutant strains that are part of the same phenotypic cluster. Remarkably, SUT125, SUT126 and SUT035 (clusters 1 and 2) dysregulate 481 coding genes (286 downregulated and 195 up-regulated) and 126 non-coding transcripts in common (Fig 4A and B). Moreover, those ncRNA deletion mutants displayed negative fitness during phenotypic analysis when growing in 22 out of the 23 media tested. To demonstrate the accuracy of our gene expression measurements, we selected a few candidate DE genes from the deletion mutants in clusters 1 and 2, and tested their mRNA levels by RT-qPCR. Among the selected genes, down-regulated and up-regulated expression fold change by qPCR were similar to the expression fold change obtained from the RNA-Seq data (S4 Fig and S2 Dataset), validating the RNA-seq data.

**Fig 4.**
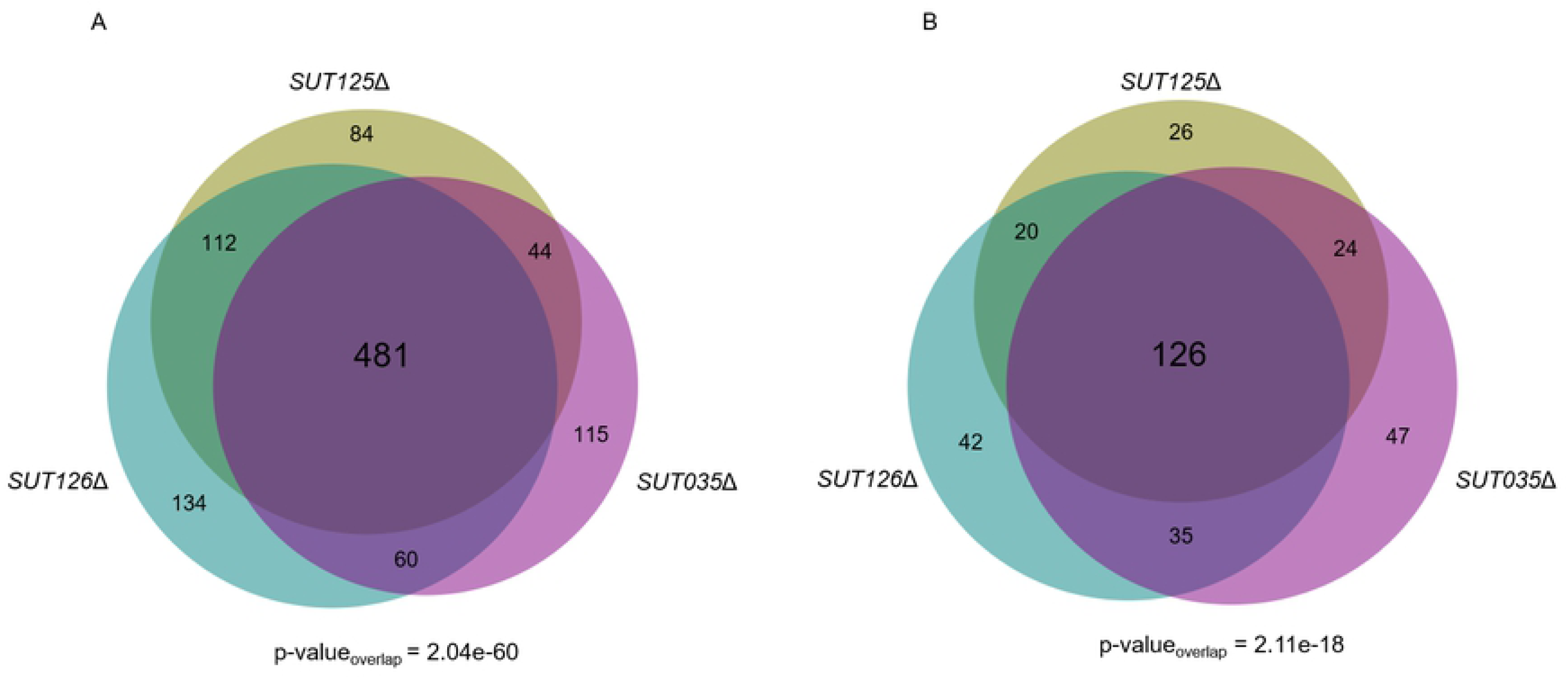
Deletion mutant strains displaying identical fitness profiles share a significant number of differentially expressed coding and non-coding transcripts. Area proportional Venn diagrams displaying the number of differentially expressed (A) Protein-coding genes (p-value= 2.04e-60 and (B) Non-coding transcripts (p-value=2.11e-18) in common between *SUT125Δ, SUT035Δ* and *SUT126Δ*. Venn diagrams were generated with BioVenn [73].

To identify the biological processes associated with the commonly misregulated genes in clusters 1 and 2, the set of DE genes was analysed for GO term enrichment and the most significant hits were selected. Genes with decreased and increased expression were associated with key mitochondrial functions such as mitochondrial electron transport and oxidation-reduction process (S5 Fig). The enriched pathways identified from KEGG and Reactome data (Holm-Bonferroni correction) were branched amino acid biosynthesis (p-value 5e-4 7 matches), aerobic respiration, electron transport chain (p-value=0.002 11 matches), mevalonate pathway (p-value= 0.004 5 matches) and TCA cycle (p-value= 0.021 9 matches), also indicating roles in mitochondrial functions. When strains in cluster 5 are included along with clusters 1 and 2, there are 96 protein-coding genes (Fig 5) and 15 non-coding genes dysregulated in common (S6 Fig). Those common genes have, in general, a concordant expression profile between each ncRNA deletion mutant strain. However, for 40% of the common genes, specifically those involved in mitochondrial function, an opposite expression trend is detected in the *SUT532*Δ strain (cluster 5) compared to *SUT125*Δ, *SUT126*Δ and *SUT035*Δ. Since the phenotypes of *SUT125*Δ, *SUT126*Δ, *SUT035*Δ and *SUT532*Δ mutants in different environmental conditions are similar; these 18 genes with different directionality of expression may either not be crucial for the observed phenotype, or specific to the mechanism of action for SUT532 in the cell (Fig 5). Due to the divergent fitness shown in glycerol for *SUT532*Δ strain we sought to elucidate if there are specific mitochondrial pathways in which SUT532 could be involved. Thus, GO of the non-common genes (321) for this ncRNA deletion mutant was also performed. Interestingly, up-regulated genes are related to the TCA cycle and aerobic respiration along with protein refolding and response to stress. Down-regulated genes are mainly involved with leucine biosynthesis biological process. (S7 Fig). Taken together, these results reveal enrichment of mitochondrial roles for SUT125, SUT126, SUT035 and SUT532 suggesting their potential function in repressing or activating mitochondrial metabolic pathways, justifying the fitness impairment of those deletion mutants when grown with non-fermentative carbon sources.

**Fig 5.**
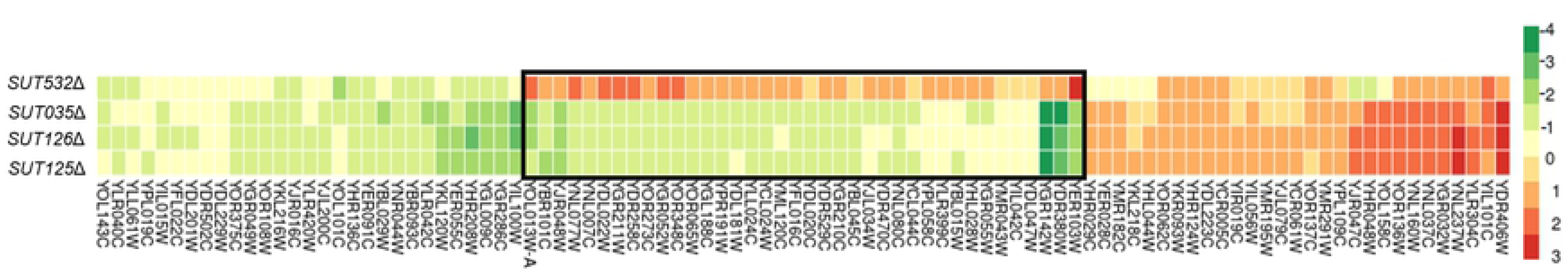
Heat map of differentially expressed genes in common between ncRNA deletion mutants with similar fitness profiles. Heat map was constructed with 96 common DE genes between *sut125Δ, SUT126Δ, SUT035Δ* and *SUT532Δ*. Colours represent the change in expression of genes, as indicated in the key on the right. DE genes in *SUT532Δ* with different transcriptional directionality from the other three ncRNA deletants are boxed.

### ncRNAs drive global transcriptome changes through transcription factors

The finding that large numbers of genes involved in the same pathways are DE in ncRNA deletion mutant strains led us to hypothesize that these ncRNAs may be acting via sequence-specific transcription factors (TFs) that regulate these groups of genes. Using the YEASTRACT database [41–44], we identified TFs that are up- or down-regulated in *SUT125*Δ, *SUT126*Δ, *SUT035*Δ, *SUT532*Δ and *CUT494/SUT053/SUT468*Δ mutants, which all show large transcriptional changes. We found that several TFs were significantly perturbed in *SUT125*Δ, *SUT126*Δ, *SUT035*Δ, *SUT532*Δ and *CUT494/SUT053/SUT468*Δ, affecting ca 16%, 19%, 20%, 13% and 5% of all annotated yeast TFs (ca. 183), respectively. The number of TFs with altered expression is significant in *CUT494/SUT530/SUT468*Δ, *SUT126*Δ, *SUT035*Δ, and *SUT532*Δ with p-values lower than 0.05 upon chi-square test (S2 Table). Several DE TFs, such as *PDR3, MOT3* and *YOX1*, were shared among *SUT125*Δ, *SUT126*Δ, *SUT035*Δ (S8 Fig). The expression changes for these three TFs were validated with *SUT126*Δ via real time PCR (S5 Fig), showing a strong agreement between the qPCR and RNA seq data.

As the most significant fitness phenotypes observed for ncRNA deletion mutant strains were in YP or YPD media supplemented with ethanol, we identified those TFs whose mis-regulation has been linked to ethanol resistance. Many ethanol-tolerance genes share a TF-binding motif recognized by Pdr1 and Pdr3 [45]. In the *S. cerevisiae* genome, 12.39% of genes are Pdr3 targets [44]. Strikingly, about 95% (*p* < 0.0001) of DE genes in *SUT126*Δ, *SUT125*Δ and *SUT035*Δ are targets of this zinc finger protein that acts predominantly as a transcriptional activator [44, 47, 48] and whose transcript levels significantly increase in the same ncRNA deletion mutant strains (S2 Dataset and S5E Fig). Furthermore, *MNS4*, which encodes a key regulator for ethanol tolerance [45,48], is up-regulated when SUT532 is deleted and down-regulated when SUT035 is deleted (S2 Dataset). Accordingly, 40.4% of dysregulated genes in the *SUT532Δ* and 37.7% in *SUT035Δ* are targets of Msn4. These data suggest that SUT125, SUT126, SUT035 and SUT532 ncRNAs are associated with mechanisms of ethanol tolerance that may involve a massive gene expression reprogramming resulting from the shift from fermentative to non-fermentative metabolism. Moreover, they imply that ncRNAs may be part of the activation or repression of metabolic pathways and regulatory networks through modulation of TFs.

To test whether the upregulation of Pdr3 target genes upon ncRNA deletion is Pdr3-dependent, we investigated the expression of previously validated Pdr3p target genes [46, 50–53] in the *SUT126Δ* background. We found that the SUT126 deletion is not sufficient to activate Pdr3 target genes *ACO1, BDH2* or *RSB1* in the absence of Pdr3 (Fig 6). These results suggest that the global effect on the transcriptome observed in the absence of SUT126 is likely driven by an effect of this ncRNA on TFs such as Pdr3. SUT126 may have a repressive effect on the promoter of *PDR3*, may destabilize the *PDR3* transcript, or, as *PDR3* is autoregulated, may bind to and interfere with the Pdr3 protein. Several ncRNAs have been reported to bind transcription factors to regulate gene expression in other organisms. For example, in mice, the long-ncRNA (lncRNA) *linc-YY1*, involved in myogenesis, has been found to interact with the TF *YY1* [24]. Similarly, *GAS5* interacts with glucocorticoid receptors, supressing their binding with glucorticoid response elements [26]. In humans, lncRNA rhabdomyosarcoma 2–associated transcript (RMST) interacts directly with Sox2, a transcription factor involved in the regulation of embryonic development [54]. Regulation of gene expression by ncRNAs acting through transcription factors might, therefore, be a conserved mechanism among eukaryotes. In this way ncRNAs could confer an extra advantage to yeast cells by modulating gene expression in response to environmental stress.

**Fig 6.**
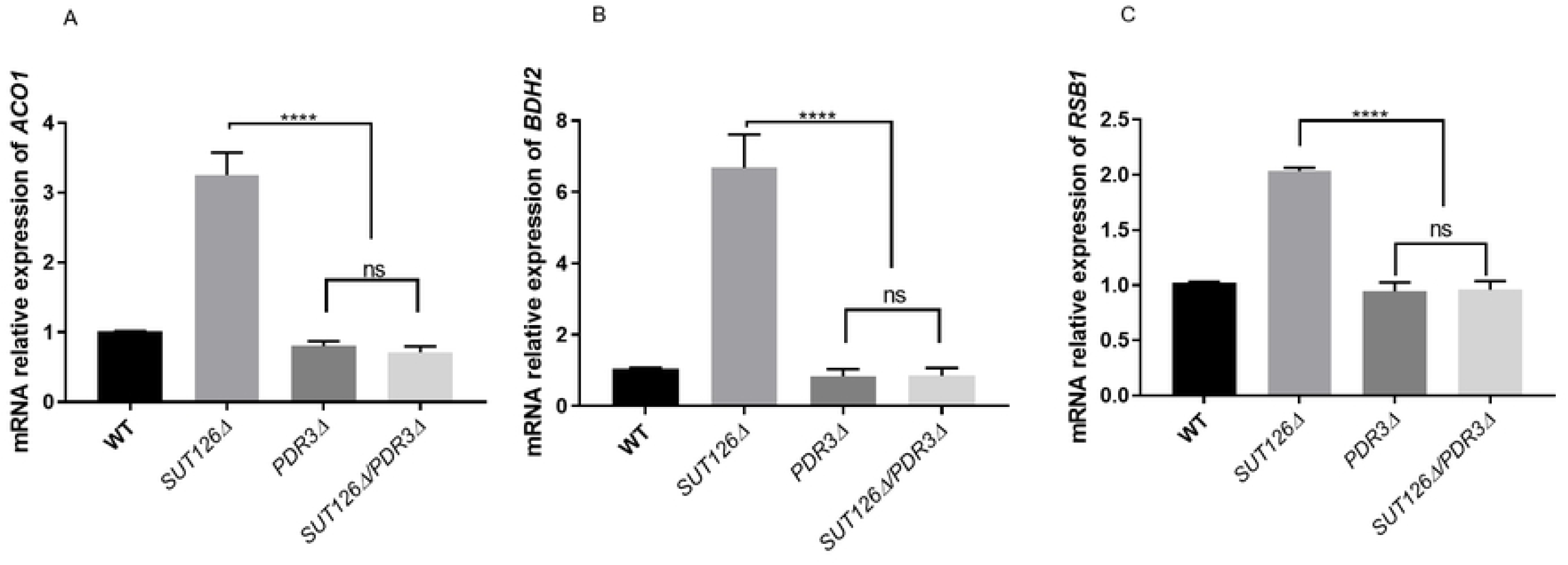
Indirect gene expression changes may be driven by SUT126 ncRNA acting through transcription factors. Relative mRNA levels of A) *ACO*1, B) *BDH2*, and C) *RSB1* analysed by RT-qPCR with *SUT126Δ/PDR3Δ* single and double mutants. The increased levels of *PDR3* targets in the *SUT126Δ* single mutant are dependent on Pdr3 (*t*-test).

### Phenotypic and transcriptional effects of the *KanMX* cassette used to generate ncRNA deletions on neighbouring genes

The *kanMX* cassette used to make the ncRNA deletion mutant strains has been suggested to affect the expression of neighbouring genes, either because of its high transcriptional level or via the generation of unexpected antisense transcripts [55–58]. We did not observe any alteration in transcript levels of neighbouring genes in the majority (13/20) of the ncRNA deletion mutant strains that we studied (Table 1) but levels of one or both neighbouring transcripts were affected in the remainder and might, therefore, contribute to the observed changes in phenotype and gene expression. For example, *SUT125Δ*, besides globally affecting the transcriptome, also has an effect on both of its neighbouring genes, *PIL1* and *PDC6*. Levels of *PIL1 mRNA* are reduced while *PDC6* transcript levels are higher in the mutant.(S3 Fig and S2 Dataset) To test whether *kanMX* cassette insertion replacing SUT125 causes the local expression changes, the mRNA levels of *PDC6* and *PIL1* were quantified and compared in three different SUT125 deletion mutant strains containing: *i*. the *kanMX* cassette in sense orientation relative to *SUT125*; *ii*. the *kanMX* cassette in antisense orientation relative to *SUT125*; *iii*. a *loxP* scar after *kanMX* excision with the Cre/loxP system (i.e. no *kanMX* cassette). The down-regulation of the expression of *PIL1* remains the same in all three mutants, ruling out a transcriptional effect of the *kanMX* cassette on *PIL1* expression (Fig 7A). *PDC6* is up-regulated in all three mutants, however the effect is stronger when the *kanMX* cassette is removed (Fig 7B). This result suggests a partial effect of the *kanMX* on *PDC6* expression, where the presence of the cassette either in sense or antisense orientation dampens the up-regulatory effect.

**Fig 7.**
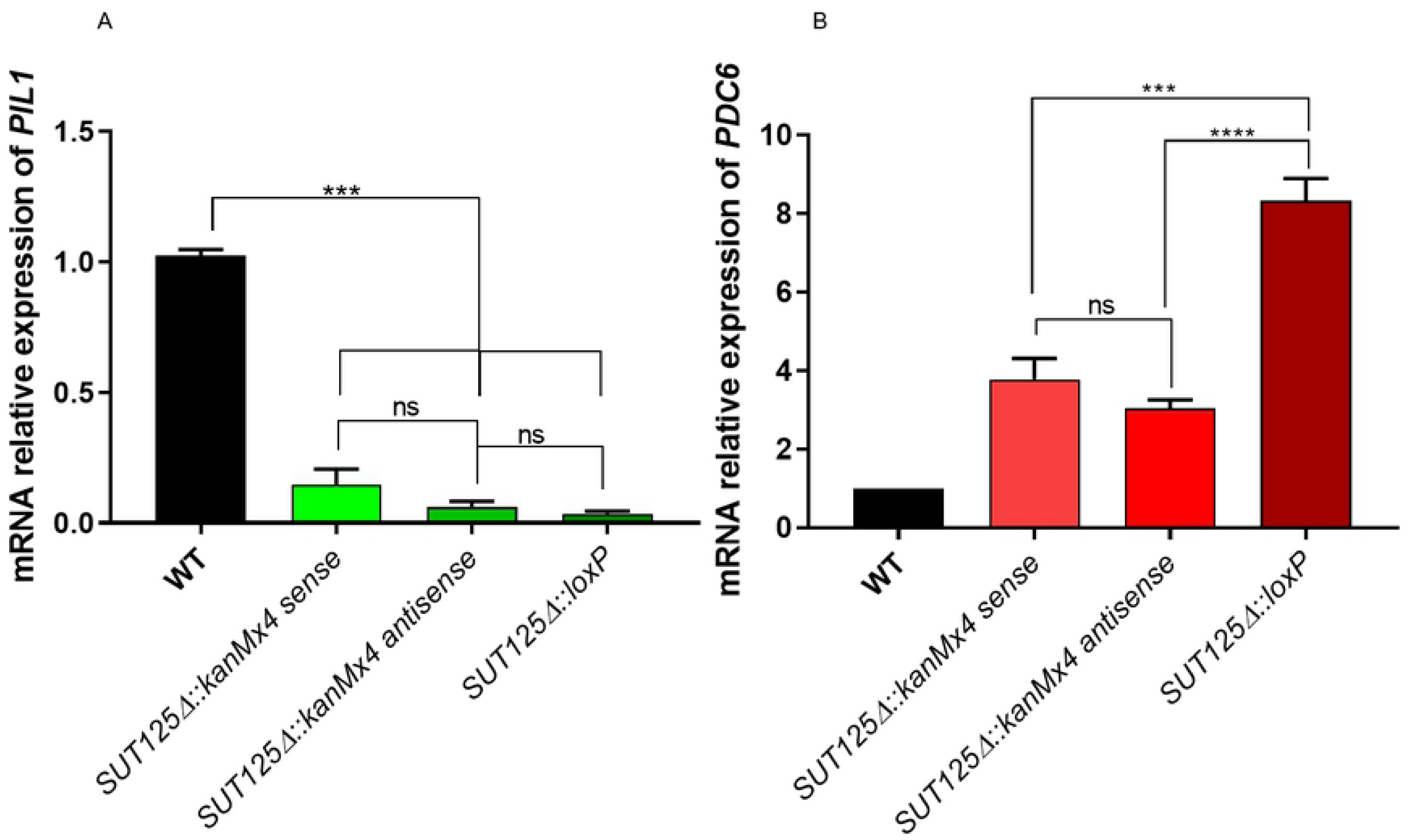
Actively transcribed *kanMX* partially decreases the regulatory effect of neighbouring genes in *SUT125* deletion strains. Transcriptional changes of SUT125 neighbouring genes (A) *PIL1* and (B) *PDC6* in *SUT125*Δ mutant strains with sense, antisense orientations (relative to SUT125) of the *kanMX* cassette, and without *kanMX* after excision with the Cre/loxP system. Relative mRNA levels were quantified by qPCR and compared by *t*-test.

To identify whether *PIL1* down-regulation and *PDC6* overexpression trigger the growth changes observed in *SUT125Δ* in medium containing ethanol we carried out spot test growth assays. *PDC6* was overexpressed from a plasmid to mimic up-regulation, and a *PIL1* deletion strain was used to mimic *PIL1* downregulation. The combined effect was scored in a *PIL1Δ* strain harbouring the *PDC6* overexpression plasmid. Presence or absence of *the kanMX* cassette reveals little to no effect on the resulting phenotype (Fig 8). Overexpression of *PDC6* in a WT background produced the same phenotype as a SUT125 deletion, while either *PIL1Δ* or *PDC6Δ* deletion did not have any effect on the phenotype (Fig 8). The concomitant effect of over-expressing of *PDC6* in *PIL1Δ* strain produced a less severe, but still comparable, phenotype to that of a SUT125 deletion. These data suggest that *PDC6* overexpression alone may account for the majority of the phenotype following *SUT125Δ* deletion.

**Fig 8.**
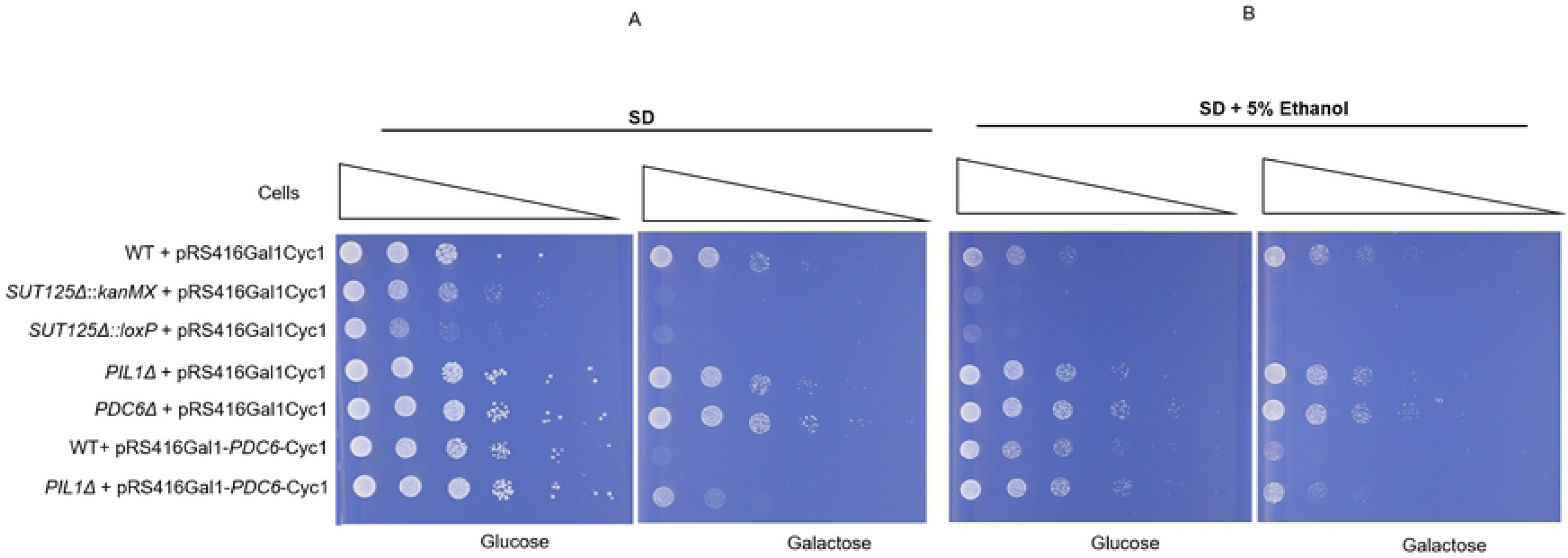
*PDC6* overexpression may explain the majority of the *SUT125Δ* phenotype. Spot test assay of: *SUT125Δ* deletion strains with and without *kanMX; PIL1Δ* deletion strain; *PDC6* overexpression strain; and *PIL1Δ* with *PDC6* overexpression plasmid plated on (A) Synthetic minimal medium lacking uracil (SD-Ura) and (B) SD-Ura + 5% ethanol, containing either 2% glucose or 2% galactose as indicated below each panel. The *PDC6* overexpression plasmid has the *PDC6* gene under control of the inducible *GAL1* promoter in the pRS416 plasmid. Wild-type and deletion strains containing the pRS416Gal1Cyc1 (empty plasmid) and the *PDC6Δ* deletion strain were included as controls.

The effect of the *kanMX* cassette on growth phenotypes was also tested in *SUT126Δ*, which has a fitness impairment, and SUT129Δ, which displays a fitness gain. Similar fitness profiles were observed regardless of the presence or absence of *kanMX* for all the ncRNA mutants (Fig 9). In addition, the effect of the *kanMX* selectable marker on transcription of non-neighbouring DE genes was tested by quantifying and comparing the mRNA levels of Pdr3p and Yox1p transcription factors in *SUT126* and *SUT125* deletion mutant strains with and without *kanMX*. No significant differences in the expression levels of *YOX1 and PDR3* were detected (S9 Fig). In summary, these data indicate that phenotypic and transcriptional changes observed in these ncRNA deletion mutants are not dependent on the presence of an actively transcribed drug resistance marker gene. Moreover, the majority of the ncRNA deletion mutants tested do not have any effect on transcript levels of neighbouring genes, suggesting a genuine effect on distant genetic loci in *trans*.

**Fig 9.**
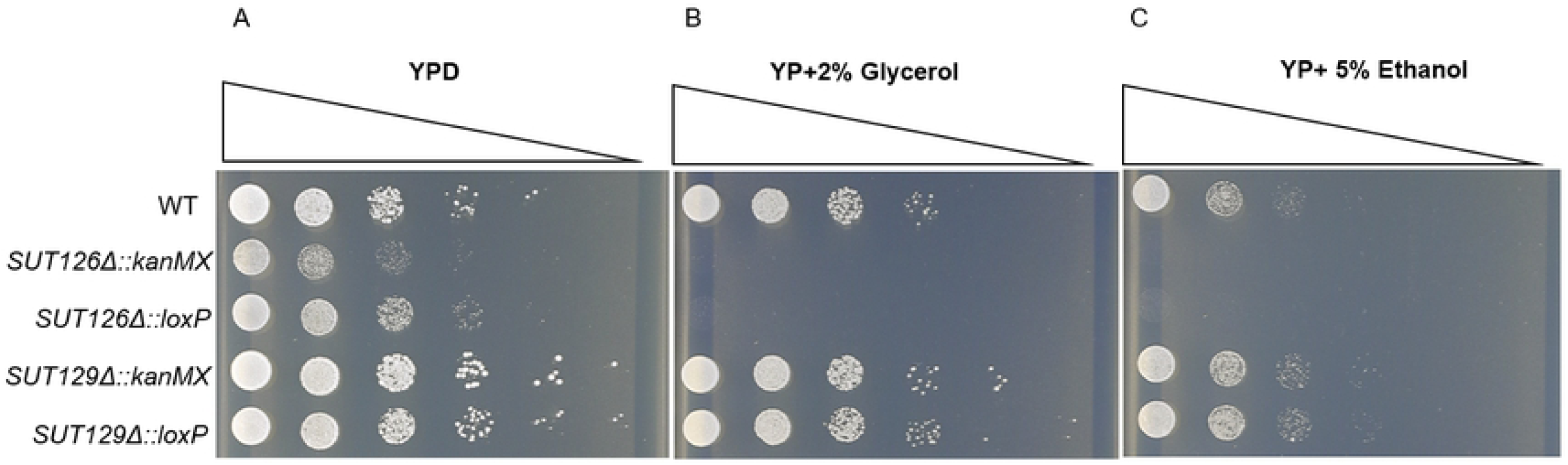
Presence or absence of *the kanMX* cassette does not affect growth phenotypes in ncRNA deletion strains. Spot test assay of: BY4741 (WT), *SUT126Δ* and *SUT129Δ* with and without *kanMX* on (A) YPD; (B) YP+ 2% Glycerol; and (C) YP+ 5% Ethanol.

### ncRNAs SUT125, SUT126, SUT035 and SUT532 act in *trans* to regulate target genes

The deletion of SUT126, SUT125, SUT035 or SUT035 led to widespread changes in the global transcription network (S2 Dataset). These ncRNAs may therefore function in *tran*s by affecting distant genes. To test this hypothesis, ectopically expressed SUT125, SUT126, SUT035 and SUT532 were assessed for their ability to rescue growth defects in the presence of 5% ethanol. Each of these SUTs was placed under control of an inducible *GAL1* promoter on a plasmid that was transformed into the respective deletion mutant. Under conditions where the *GAL1* promoter is repressed (glucose) there ware no differences in growth between deletion strains carrying the *GAL1-SUT* plasmid or an empty version of this plasmid. However, when *GAL1*-driven expression was induced (galactose), all four SUTs were able to rescue the growth defect (Fig 10). These results suggest that SUT126, SUT126, SUT035 and SUT532 can act *in trans*, which may underlie the altered regulation of large numbers of genes in these mutants. There are only a handful of examples of *trans* acting ncRNAs in yeast but a CUT that affects gene regulation, CUT281, can act both in *cis* and *trans* to repress the *PHO84* gene [6, 36], while SUT457 can act in *trans* to rescue the phenotype of telomeric overhang accumulation observed in *SUT457Δ* cells [9].

**Fig 10.**
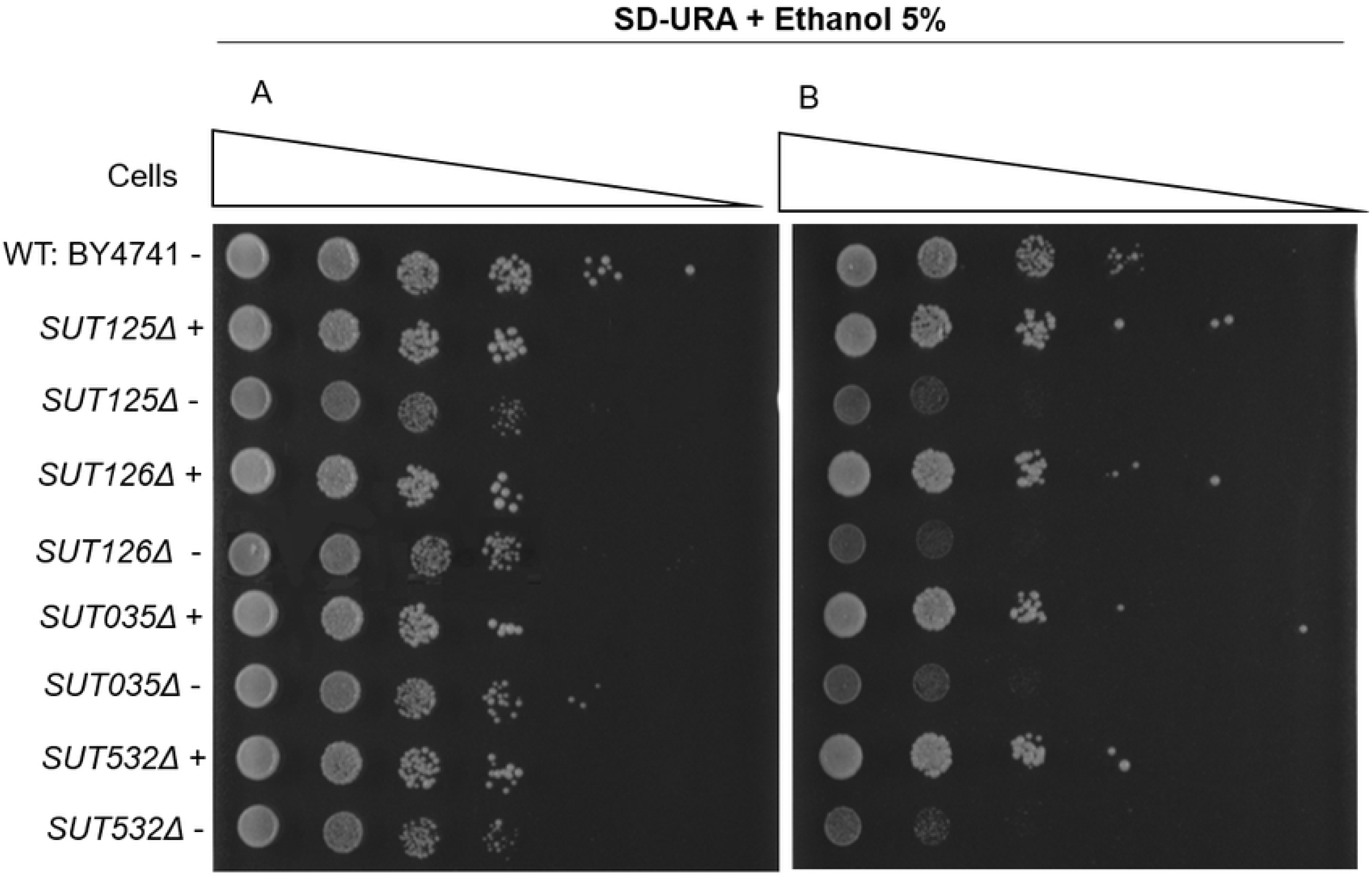
SUTs whose deletion results in wide-spread transcriptional changes can rescue growth phenotypes in *trans*. Rescue spot test analysis of growth phenotypes of ncRNA deletion mutant strains containing the indicated SUT under control of the *GAL1* promoter, spotted onto SD + 5% ethanol, containing either 2% glucose (A) or 2% galactose (B). +: pRS416 with the respective SUT; - : empty plasmid.

## Conclusion

Large-scale phenotypic projects using deletion mutant collections have proven to be an invaluable tool for linking genes to their function [59–62]. Here we used 372 haploid strains from the ncRNA deletion collection [10] to identify deletions that are responsible for phenotypic changes in 23 environmental conditions. The fitness data obtained has been integrated into the Yeast ncRNA Analysis (YNCA) database (http://sgjlab.org:3838/ynca/) [10]. Based on the phenotypic screening data, we further analysed 20 ncRNA deletion mutants at the transcriptome level. ncRNA deletion mutants that were phenotypically impaired also triggered significant changes in the gene regulatory network. By analysing the expression data, we identified specific pathways where these SUTs and CUTs were functioning, such as mitochondrial function and respiration, ethanol tolerance, rRNA processing, plasma-membrane fluidity and sterol biosynthesis. In the *SUT126*Δ strain, we showed that the large transcriptional changes are due to the altered expression of TFs rather than the direct effect of the lncRNA deletion. These results indicate that ncRNAs are likely to be involved in fine tuning expression by regulating the expression of TFs.

Gene regulation driven by ncRNAs through TFs may be a conserved mechanism amongst eukaryotes. Examples of ncRNAs enhancing the loading of TFs at their target promoters or acting as a binding competitors for DNA/RNA binding proteins in fission yeast, mouse and human cells are increasing [23–26]. In fact, most ncRNAs are transcribed near regulatory units for transcription such as promoters or enhancers [17, 18, 63], which may be an indication that associates them with biological function and mechanism.

We discovered that SUT125, SUT126, SUT532 and SUT035 act in *trans* since their functions can be rescued ectopically. Strikingly, these ncRNAs originate from intergenic regions that do not overlap with any open reading frame, bearing out the possibility that their functionality may be linked with their potential to form accessible structural domains able to bind to DNA, RNA or proteins [64–66]

Such ncRNA mediated regulation is cost-effective compared to the classical regulation via TFs as the fast production of RNAs compared to proteins facilitates quick genetic responses to environmental stimuli.

## Materials and methods

### Yeast strains, growth conditions and plasmids

A list of *Saccharomyces cerevisiae* strains and plasmids is provided in S3 Table. For strain maintenance and construction, strains were grown at 30°C under standard conditions. ncRNA single deletion strains used this study were taken from the ncRNA deletion collection created by Parker, et al [10,67]. Deletion mutants were maintained on Yeast extract Peptone Dextrose Agar (YPDA) containing 200 µg/mL G418. Double deletion mutant strains were constructed by substituting the candidate *SUT* locus with the *natNT2* cassette and were maintained on YPDA containing 100 µg/mL clonNAT.

For construction of strains ectopically expressing particular SUTs, isogenic wild-type and ncRNA deletion mutant strains cells were transformed with pRS416-Gal1-Cyc1 overexpression plasmid containing the ncRNA of interest. Resulting strains were maintained in a synthetic minimal media lacking uracil (SD-Ura: 1X Yeast Nitrogen Base (YNB) (Formedium); 1X Complete Supplement Mixture (CSM) – Ura (Formedium); 2% (w/v) glucose). For phenotypic rescue studies, strains were grown to an optical density at 600 nm (OD600) of 0.5 in YP (1% yeast extract, 2% peptone) medium supplemented with 2% raffinose (YPRaf) at 30°C and induced with YP medium containing 2% galactose (YPGal) for 2 hours before being harvested for spot test assays.

### Cre recombinase-mediated marker excision in *Saccharomyces cerevisiae*

SUT deletion strains containing loxP sites flanking the *kanMx* cassette were transformed with pSH-bler plasmid DNA, and grown on YPDA containing 10 µg/mL phleomycin. To excise the cassette, cells harboring pSH-bler were grown overnight in YPRaf medium, re-suspended in 10 ml YPGal medium to an OD600 of 0.3 and incubated at 30°C for 3 h. The culture was diluted and plated out on YPDA. The resulting colonies were replica-plated on YPDA containing 200 µg/mL G418 to confirm the marker loss and YPDA with 10 µg/mL phleomycin to confirm the plasmid loss. The marker loss was also verified by colony PCR.

### Phenotypic analysis on solid and liquid media

Two biological and four technical replicates of the haploid deletion mutant strains were arrayed in 384 well microtitre plates. Using a Singer Rotor HDA, the 384 well cell cultures were stamped onto YPDA plates and replica plated onto 23 different environmental conditions and incubated at a particular temperature. A full list of the media and temperatures used in this study are listed in S5 Table. Plates were imaged at 24, 48 and 72 hours using a Bio-Rad Gel Doc XR system and images were processed using SGAtools [68]. The average pixel count for the replicates of each strain were then normalized to the appropriate plate wild-type value then mean, standard deviation and p-values were calculated assuming a normal distribution of values. Strains with similar growth in different media were grouped into specific clusters.

For liquid fitness assays, cells were grown at 30°C from an OD600 nm of 0.1, and growth measurements at OD595nm were recorded using a BMG FLUOstar OPTIMA Microplate Reader. The readings were taken every 5 minutes as previously described by Naseeb and Delneri [69] for up to 55 hours incubation time. Three technical replicates of three independent biological samples were used for each deletion mutant and wild-type strain. Graphs and growth parameters were produced using the *grofit* package of the *R* program.

For spot test assays, cultures were grown overnight before being serially diluted 1:10 and spotted onto agar plates.

### Total RNA extraction and quantitative RT-PCR

Total RNA was isolated from 1×10^7^ cells using the RNeasy Mini Kit (QIAGEN, Germany) following the protocol for enzymatic digestion of cell wall followed by lysis of spheroplasts. To eliminate genomic DNA contamination, an additional DNAse treatment was performed with RNAse-free DNase set (QIAGEN, Germany) following the manufacturer’s protocol. The RNA extracted was quantified using a NanoDrop LiTE Spectrophotometer (THERMO SCIENTIFIC, United States). Two micrograms of total RNA were reverse transcribed into cDNA using SuperScript III Reverse Transcriptase (Invitrogen, UK) according to the manufacturer’s protocol. Optimized qPCR reactions contained 2ng/µl of cDNA, 3pmol each primer and 5 µl of iTAq Universal SYBR Green super Mix 2X in a final volume of 10 µl. Reactions were cycled on a Roche Light Cycler real time System for 35 cycles of: 15 seconds at 95°C; 30 seconds at 57°C ;and 30 seconds at 72°C. Three biological replicates and three technical replicates per sample were used in each experiment, and all runs included a no template control, and a control lacking reverse transcriptase. The relative expression of each gene was estimated using the Ct values relative to those of *ACT1*. Primers were designed to produce an amplicon of 80-150bp (Sequences given in S4 Table).

### Illumina HiSeq library preparation and sequencing

Libraries were prepared from total RNA using the TruSeq Stranded mRNA Library Prep Kit (Illumina,Inc) according to the manufacturer’s instructions. Sequencing was performed on an Illumina HiSeq4000 instrument. Sequences corresponding to protein-coding genes were mapped to sacCer3, while CUT and SUT sequences were mapped using the genomic coordiates provided by Xu et al [17]. Mapping was performed using STAR [70]. Differential gene expression analysis was based on the negative binomial distribution (DESeq2) [71]. Genes with a statistically significant difference in expression from wild-type, as indicated by a q-value below 0.1, and greater than 1.5 fold change in expression, were included in the final list of differentially expressed genes.

### Bioinfomatic and statistical analyses

Differentially expressed genes were listed and grouped as up- or down-regulated. Enriched GO terms and pathways were identified using YeastMine, with the Helmed-Bonferroni correction used to calculate adjusted *p*-values [72]. The Yeast Search for Transcriptional Regulators And Consensus Tracking (YEASTRACT) [44] database was used to look for transcription factors and their target genes.

Statistical tests were performed using Welch two sample t-test and multiple comparisons were analysed using ANOVA followed by Dunnett’s test. Error bars denote standard deviations except where noted and *p*-values are indicated on Figs as: .*\* p < 0.05 ** p < 0.01 ***p < 0.001 ****p <0.0001;* ns = no significant change.

## Author Contributions

DD, CBM and ROK conceived the study; DD, CBM and LNB designed the experiments; LNB, SP and MF performed the experiments, PW and ST contributed to the initial assembly and normalisaiton of RNAseq data; LNB, CBM and DD analysed data; LNB, CBM and DD wrote the paper with the input of SP and ROK.

## Acknowledgements

We thank the Genomic Technologies and Bioinformatics Facilities in the Faculty of Biology, Medicine and Health for their contribution in the acquisition and mapping of RNA-Seq data.

## Supporting information

### S1 Dataset

This file contains the solid fitness data of the 372 mutant strains tested in 23 different conditions. It contains a summary of the clusters and the p-value per strain per condition

### S2 Dataset

This file contains the RNA-seq data divided by mutant, containing the list of significant DE genes per mutant strain. Tables are divided by protein-coding genes and non-coding transcripts

**S1 Fig. Solid fitness of heterozygous deletions of essential ncRNAs, SUT075 and snR30**

Bar charts displays the colony size of *SUT075Δ* and *snR30Δ* deletion strains when growing in (A) YPD and (B) YPD supplemented with 10% ethanol.

**S2 Fig. Gene ontology for biological process enriched in DE genes in common between snR30 and SUT075**. Bar chart displaying the 20 first significantly enriched GO terms. The negative logarithm of the adjusted p-value (base 10) after Holm-Bonferroni correction is represented on the x-axis. The figure was created using the DE genes in common for SUT075 and snR30 deletion mutants. n=1836.

**S3 Fig. Histogram of GO terms from DE genes in *CUT494/SUT530/SUT468Δ* strain**. Representative GO terms for biological processes for up-regulated (red) and down-regulated (green) genes in the *CUT494/SUT530/SUT468Δ* strain. Holm-Bonferroni p-value cutoff < 0.05; y –axis displays GO terms, x-axis shows the p-value that was transformed to –log10. The figure was created using the DE genes. n=137.

**S4 Fig. Validation of DE genes obtained during RNA-seq by qPCRs**. Relative mRNA levels of (A) *PDR3* and (B) *YOX1* in *SUT035*Δ strain, (C) *PDC6* and (D) *PIL1* in *SUT125*Δ and the TFs (E) *PDR3*, (F) Y*OX1* and (G) *MOT3* in *SUT126*Δ strain analyzed by RT-qPCR. Relative mRNA levels were quantified by qPCR and compared by *t*-test.

**S5 Fig. SUT125, SUT126 and SUT035 reveal an important role in mitochondrial processes**. Gene Ontology of biological processes inferred from dysregulated coding targets in common in *SUT125*Δ, *SUT126*Δ and *SUT035*Δ deletion strains. Significantly first enriched GO terms for biological processes (Holm-Bonferroni adjusted p-value <0.05) are listed on the y-axis, and the negative log of the adjusted p-value (base 10) is represented on the x-axis. The figure was created using the DE genes in common for SUT125, SUT126 and SUT035 n=481.

**S6 Fig. Gene Ontology of biological processes inferred from DE protein coding genes in S*UT532Δ* deletion mutant strain**. Significantly enriched representative GO terms for biological processes for up-regulated (red, n=172) and down-regulated (green, n=236) in *SUT532Δ* deletion strain. P-value was calculated using Holm-Bonferroni correction. Representative GO terms are listed on the y-axis, and the negative log of the adjusted p-value (base 10) is represented on the x-axis.

**S7 Fig. Area proportional Venn diagram of DE transcripts between cluster 1, 2 and 5**. Number of (A) Protein coding genes (96) and (B) Non-coding transcripts (15) in common dysregulated among deletion strains in cluster 1 (*SUT125Δ, SUT035Δ*), 2 (*SUT126Δ*) and 5 (*SUT532Δ*). Venn diagram generated using Eulerr [74]

**S8 Fig. Venn diagram representing TFs in common between phenotypic related ncRNA deletion mutants with significant impact on the genome**. Area proportional Venn diagram generated by BioVenn [73] using the number of TFs dysregulated in deletion strains in cluster 1 (*SUT125*Δ, *SUT035*Δ) and 2 (*SUT126*Δ). The overlapping (23 TFs) is shown in a dark green colour.

**S9 Fig. Altered expression levels of target genes in ncRNA deletion mutant strains are independent of *kanMX* marker**. Relative mRNA levels of the transcriptional repressor (A) *YOX1* and the transcriptional activator (B) *PDR3* in *SUT125*Δ and *SUT126*Δ deletion mutant strains with and without *kanMX*. The *kanMX* cassette does not influence genes located distantly from the SUT disruption. Relative mRNA levels were quantified by qPCR and compared by *t*-test.

**S1 Table**. Characteristic parameters of growth curves of deletion mutant strains assessed in liquid media. Tables show mean values normalized with wild type, standard deviation (SD), adjusted p-value and significance per parameter.

**S2 Table**. List of transcription factors DE in mutant strains.

**S3 Table**. List of yeast strains and plasmids used in this study

**S4 Table**. List of primers for Quantitative real time PCR (qPCR) used in this study

**S5 Table**. List of media condition used for solid fitness analysis for the haploid ncRNA deletion collection.

